# Haplotype-resolved chromosome-level genome assemblies of four *Diamesa* species reveal the genetic basis of cold tolerance and high-altitude adaptations in arctic chironomids

**DOI:** 10.1101/2025.07.30.667619

**Authors:** Sarah L.F. Martin, Renato La Torre, Bram Danneels, Ave Tooming-Klunderud, Morten Skage, Spyridon Kollias, Ole Kristian Tørresen, Mohsen Falahati Anbaran, Elisabeth Stur, Kjetill S. Jakobsen, Michael D. Martin, Torbjørn Ekrem

## Abstract

Arctic and alpine insects face extreme environmental stressors, yet the genomic basis of their adaptation remains poorly understood. Here, we present the first haplotype-resolved, chromosome-level genomes for four species of *Diamesa* (Diptera: Chironomidae), a genus of cold-adapted midges inhabiting glacial and high-altitude freshwater ecosystems. Using PacBio HiFi sequencing and Hi-C scaffolding, we assembled high-quality genomes with chromosome-level resolution and high k-mer completeness. Phylogenomic analyses support Diamesinae as sister to other Chironomidae except Podonominae, and genomic comparisons provide evidence for introgression between the evolutionary distinct *D. hyperborea* and *D. tonsa*. Comparative genomic analyses across 20 Diptera species revealed significant gene family contractions in *Diamesa* associated with oxygen transport and metabolism, suggesting adaptations to high-altitude, low-oxygen environments. Conversely, expansions were detected in histone-related and Toll-like receptor gene families, likely enhancing chromatin remodeling and immune regulation under cold stress. A single gene family encoding glucose dehydrogenase was significantly expanded across all cold-adapted species studied, implicating its role in cryoprotectant synthesis and oxidative stress mitigation. Notably, *Diamesa* species exhibit the largest gene family contraction at any node, with minimal overlap in expansions with other cold-adapted Diptera, indicating lineage-specific adaptation. Our findings support the hypothesis that genome size condensation and selective gene family changes underpin survival in cold environments. These genome assemblies represent a valuable resource for investigating adaptation, speciation, and conservation in cold-specialist insects. Future work integrating gene expression and population genomics will further illuminate the evolutionary resilience of *Diamesa* in a warming world.

## INTRODUCTION

Understanding the effects of climate change on insect populations is of fundamental importance for management and conservation of freshwater and terrestrial ecosystems [1]. While dispersal to habitable areas is a response option for lowland and temperate taxa, high altitude or arctic species have nowhere to run to as temperatures and other climatic variables change to levels beyond those critical for their survival [2]. Remaining then, is the species’ ability to adapt to the changing environment, either through altered behaviour, change in phenology or physical appearance [3]. The adaptive potential (i.e. the capacity to respond to changing selection pressures) of an insect is dependent on multiple factors including genomic mechanisms [4]. Thus, our ability to understand an organism’s capacity to cope with environmental change should include knowledge of its genome, perhaps especially of non-model organisms with adaptations to extreme environments [5].

Midges of the family Chironomidae (Diptera) are among the most abundant and species-rich aquatic insects worldwide [6]. The family is represented in all continents and biogeographic regions, including the Antarctic mainland [7], and is one of few insect groups in which multiple evolutionary lineages have adapted to life in the marine environment [6]. While the immature stages of the majority of species are aquatic, terrestrial and semi-terrestrial species are also common[8]. Some species have adapted to life in extreme environments and are capable of enduring desiccation [9], heavy pollution [10], low pH [11], high salinity [12] and low temperatures [13]. Species of the subfamily Diamesinae typically have immatures associated with cold, flowing waters or nutrient poor lakes [14], with the genus *Diamesa* colonizing such habitats mainly in the northern hemisphere and in some regions being valuable bioindicators of cold mountain waters [13]. A better understanding of the genomic architecture in *Diamesa* species could therefore help identify regions associated with adaptation to cold temperatures in these insects. Moreover, as the genus *Diamesa* has seen a rather recent radiation in the Neogene period [13,14], with reported lower evolutionary rates of mitochondrial protein-coding genes [15] and suspected hybridization [16], it is of interest to explore the genomic divergence between closely related species.

Advances in long-read sequencing technologies, such as PacBio and Oxford Nanopore, in combination with Hi-C scaffolding, have made it possible to generate chromosome-level genome assemblies even for non-model organisms with small genomes, such as *Diamesa* species. These modern approaches provide vastly improved contiguity and completeness compared to earlier short-read assemblies, such as the genome of *Belgica antarctica* [17], which was highly fragmented and lacked chromosomal resolution. Importantly, global initiatives like the Earth BioGenome Project are accelerating the production of high-quality reference genomes across the tree of life, including ecologically important but historically understudied taxa. By providing standardized protocols and infrastructure, these efforts are enabling comprehensive biodiversity genomics and helping to close longstanding taxonomic and genomic gaps.

The genome of the Antarctic midge *Belgica antarctica* was the first chironomid genome to be published [17]. At the time, it was the smallest insect genome sequenced (99 Mbp), which was featured as a likely adaptation to an extreme environment. Sequenced genomes from other midges in Chironomidae as well as the sister family Ceratopogonidae indicate, however, that the small genome size is a plesiomorphic trait for the family [18]. Over the last decade, there have been published chromosome-level genomes of ten species in four subfamilies: *Paroclus steinenii* (Podonominae), *Clunio marinus, Smittia aterrima* and *Smittia pratorum* (Orthocladiinae), *Chironomus riparius, Chironomus tentans, Polypedilum pembai, Polypedilum vanderplanki* and *Tanytarsus gracilentus* (Chironominae), and *Propsilocerus akamusi* (Prodiamesinae) (Table 1). Thus, after the recent establishment of the Protanypodinae [14], there are eight subfamilies within the Chironomidae that lack published genomes. While some of the previously published studies on chironomid genomes are largely descriptive (e.g. [19,20]), others discuss genomic mechanisms underlying tolerance to heavy metal exposure [10], anhydrobiosis [21], low temperatures [17], or generally stressful environments [18]. For the marine midge *Clunio marinus*, whose reproduction is timed with tide, Kaiser et al. [22] used the genomes from five geographically different lineages to map loci for circalunar and circadian chronotypes. It is worth mentioning that closer examination of the COI barcodes from the populations used to generate the *Smittia* genomes [19] indicate that these do not belong to the species assigned in the publication, but other members of the same genus.

**Table 1.** Available whole genomes in Chironomidae.

| Species | Subfamily | Total length (Mbp) | BUSCO completeness (%) | Accession number | Reference |
| --- | --- | --- | --- | --- | --- |
| *Parochlus steinenii* | Podonominae | 143.57 | 92.27 | GCA_038502155.1 | [18,23] |
| *Belgica antarctica* | Orthocladiinae | 89.58 | 91.72 | GCA_000775305.1 | [17] |
| *Clunio marinus* | Orthocladiinae | 85.49 | 93.18 | GCA_900005825.1 | [22] |
| *Smittia aterrima* * | Orthocladiinae | 78.45 | 98.5 | GCA_033063855.1 | [19] |
| *Smittia pratorum* * | Orthocladiinae | 71.56 | 97.8 | GCA_033064975.1 | [19] |
| *Chironomus riparius* | Chironominae | 191.84 | 92.60 | GCA_917627325.3 | [20] |
| *Chironomus tentans* | Chironominae | 213.46 | 89.35 | GCA_000786525.1 | [24] |
| *Polypedium pembai* | Chironominae | 122.92 | 91.90 | GCA_014622435.1 | [25] |
| *Polypedium vanderplanki* | Chironominae | 118.97 | 93.18 | GCA_018290095.1 | [21] |
| *Tanytarsus gracilentus* | Chironominae | 91.83 | 91.57 | GCA_038502055.1 | [18] |
| *Prosilocerus akamusi* | Protanypodinae | 85.84 | 92.79 | GCA_018397935.1 | [26] |

\*Comparison of COI barcodes indicate that these genomes belong to other species in the genus *Smittia*.

Here we generate high-quality haplotype-resolved chromosome-level assemblies for four species of *Diamesa* (subfamily Diamesinae), the first genomes generated for this subfamily using long read sequencing (PacBio HiFi) and long range chromosomal contact maps (Hi-C). We use these genomes to investigate the evolutionary mechanisms employed by these species to overcome extreme environments, with a particular focus on cold and high-altitude tolerance.

## MATERIAL AND METHODS

### Biological sample collection and identification

Fieldwork specifically for this project was conducted at 1100 m elevation in the Rondane National Park, Norway in July 2022, where previous studies had documented 12 different species of *Diamesa*. Adult specimens were caught by sweeping vegetation near the stream Vidjedalsbekken and live specimens individually identified to genus level in the field by isolating them in 2 ml glass vials. One leg from each selected specimen was separated from the main body and preserved in ethanol, while the remaining body was snap-frozen in liquid nitrogen, labelled with consecutive numbers to ensure future association of body parts, and stored at -80℃ until ready for DNA extraction.

All *Diamesa* specimens were DNA barcoded using the following procedure. DNA from ethanol preserved legs was extracted using Qiagen DNeasy Blood and Tissue kit following the standard protocol, except elution was done twice with 50 µL buffer, using the elute from the first round in the second to increase the final DNA concentration. PCR of the cytochrome c oxidase subunit 1 (COI) barcode fragment was conducted using the primers LCO1490 and HCO2198 [27] and Qiagen Multiplex PCR Kit and a temperature profile of 1 cycle of 95℃ for 5 minutes initial denaturation, 5 cycles of 94℃ for 30s denaturation, 45℃ for 30s annealing and 72℃ for 60s for extension followed by 35 cycles of 94℃ for 30s, 51℃ for 30s and 72℃ for 60s, and 1 cycle of 72℃ for 5 minutes final extension. PCR products were cleaned using ExoSAP-IT and sequenced in the reverse direction at Eurofins Genomics (Germany) using Sanger sequencing and BigDye termination. Sequences were trimmed for uncertain base calls at each end and identified against the entries in the Barcode of Life Data Systems (BOLD [28]) with special reference to previous records from the same locality. All sequences and metadata are available in BOLD under the dataset “DS-DIANOR1 Norwegian Diamesa for genomics”, DOI: dx.doi.org/10.5883/DS-DIANOR1. Based on morphological identification (see Figure 1 for example) using available literature [29–31] and the results from DNA barcoding, male imagines of *Diamesa hyperborea, D. tonsa* and *D. serratosioi*, and one female of *D. lindrothi* were selected for individual high molecular weight (HMW) DNA extraction. Prior to placement of snap-frozen specimens in extraction buffer, wings, antennae and hypopygium (e.g. Figure 1) of each specimen were dissected off and mounted in Euparal on slides for permanent storage as vouchers in the NTNU University Museum Natural History Collections (NTNU-VM, Supplementary Table 1).

**Figure 1.**
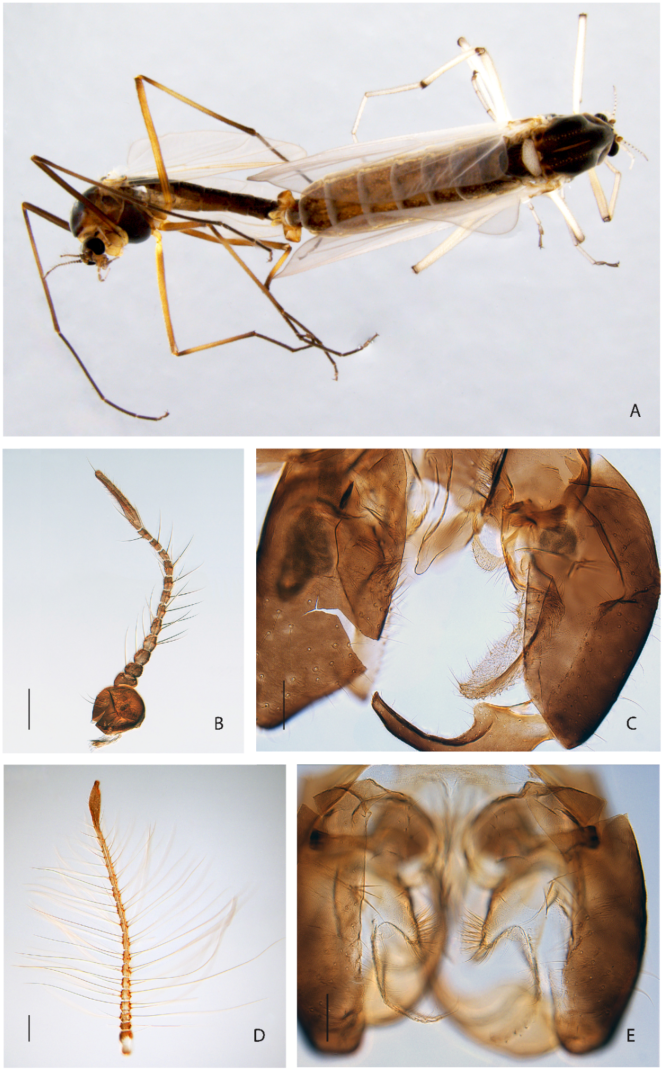
Morphological characteristics of *Diamesa hyperborea* (A-C) and *Diamesa tonsa* (D-E). A, male and female adult in copula; B & D, adult male antenna; C & E, male hypopygium. Scale bar in B & D = 100 μm, scale bar in C & E = 75 μm. Photos: Elisabeth Stur & Torbjørn Ekrem.

Additional specimens were required for Hi-C analyses (locations in Figure 1). For this we used individual male adults of *Diamesa hyperborea, D. tonsa, D. serratosioi* and *D. lindrothi* collected in 2008 and 2014 preserved in 96% ethanol and stored at 4-5℃ (details in Supplementary Table 1). These were also collected at or near the field locality, except *D. tonsa* which was collected at River Gaula near Kvål in Melhus kommune, Trøndelag, Norway (Supplementary Table 1).

### DNA extraction

After collection, specimens were stored at -80℃. Prior to DNA extraction, the specimens were taken from -80℃ for brief microscopic examination in which genitalia were removed for voucher collection. HMW DNA was extracted using the MagAttract HMW DNA kit (Qiagen) following the manufacturer’s protocol. The individual specimens were manually homogenized using a sterile pestle in the ATL buffer. The final elution was in 50 ul Buffer AE.

### Library preparation and sequencing

Before PacBio library preparation, gDNA was purified an additional time using AMPure PB beads (1:1 ratio). For two of the species, short fragment removal was performed using 0.5x AMPure beads (*Diamesa serratosioi*) and 35% diluted AMPure PB beads (*D. hyperborea*). For all four species, 16-25 ng DNA was sheared into an average fragment size of 10-15 kbp large fragments using g-tubes (Covaris). Libraries were prepared following the PacBio protocol for low input DNA: Procedure & Checklist — Preparing HiFi SMRTbell Libraries from Ultra-Low DNA Input. Libraries were size selected using 35% diluted AMPure PB beads and following Procedure & Checklist – Using AMPure® PB Beads for Size-Selection. Final libraries were pooled before sequencing on the PacBio Sequel IIe instrument (Pacific Biosciences Inc). The libraries were sequenced on one 8M SMRT cell using the Sequel II Binding kit 2.2 and Sequencing chemistry v2.0. To increase the amount of data for *Diamesa lindrothi*, the library was sequenced on approximately 5% of 25M SMRT cell on Revio instrument (also Pacbio) using Revio polymerase and sequencing chemistry. PacBio library prep and sequencing was performed by the Norwegian Sequencing Centre at University of Oslo.

For all species, the whole organism was used for generating Hi-C data. Hi-C libraries were prepared using the Arima High Coverage Hi-C kit, following the manufacturer’s recommendations for low input samples and the user guide for animal tissues (document no A160162 v01). For three of the species, one individual stored in EtOH was used as input, while for *Diamesa lindrothi*, two individuals stored in EtOH were used as input to Hi-C library prep. Final libraries were quantified using the Kapa Library quantification kit for Illumina (Roche Inc.) and pooled with other libraries before sequencing on the Illumina NovaSeq X with 2*150 bp paired end mode (lllumina Inc) at the Norwegian Sequencing Centre.

### Genome assembly

A full list of relevant software tools and versions is presented in Supplementary Table 2. We assembled the species using a pre-release of the EBP-Nor genome assembly pipeline (https://github.com/ebp-nor/GenomeAssembly). KMC [32] was used to count k-mers of size 32 in the PacBio HiFi reads, excluding k-mers occurring more than 10,000 times. GenomeScope [33] was run on the *k*-mer histogram output from KMC to estimate genome size, heterozygosity and repetitiveness while ploidy level was investigated using Smudgeplot [33]. HiFiAdapterFilt [34] was applied on the HiFi reads to remove possible remnant PacBio adapter sequences. The filtered HiFi reads were assembled using hifiasm [35] with Hi-C integration resulting in a pair of haplotype-resolved assemblies, pseudo-haplotype one (hap1) and pseudo-haplotype two (hap2). Unique k-mers in each assembly/pseudo-haplotype were identified using meryl [36] and used to create two sets of Hi-C reads, one without any k-mers occurring uniquely in hap1 and the other without k-mers occurring uniquely in hap2. K-mer filtered Hi-C reads were aligned to each scaffolded assembly using BWA-MEM [37] with -5SPM options. The alignments were sorted based on name using samtools [38] before applying samtools fixmate to remove unmapped reads and secondary alignments and to add mate score, and samtools markdup to remove duplicates. The resulting BAM files were used to scaffold the two assemblies using YaHS [39] with default options. FCS-GX [40] was used to search for contamination. Contaminated sequences were removed. The mitochondrion was assembled from PacBio HiFi reads using Oatk [41] and a minimum synmcer coverage threshold value of either 150 or 100. All the evaluation tools have been implemented in a pipeline, similar to assembly and annotation (https://github.com/ebp-nor/GenomeEvaluation). Merqury [36] was used to assess the completeness and quality of the genome assemblies by comparing to the k-mer content of the Hi-C reads. BUSCO [42] was used to assess the completeness of the genome assemblies by comparing against the expected gene content in the insecta lineage. Gfastats [43] was used to output different assembly statistics of the assemblies. The assemblies were manually curated using PretextView and Rapid curation 2.0. Chromosomes were identified by inspecting the Hi-C contact map in PretextView. BlobToolKit and BlobTools2 [44], in addition to blobtk, were used to visualize assembly statistics. To generate the Hi-C contact map image, the Hi-C reads were mapped to the assemblies using BWA-MEM [37] using the same approach as above, before PretextMap was used to create a contact map which was visualized using PretextSnapshot. The mitochondrial genome was recovered for all species.

### Genome annotation

We annotated the genome assemblies using a pre-release version of the EBP-Nor genome annotation pipeline (https://github.com/ebp-nor/GenomeAnnotation). First, AGAT (https://zenodo.org/record/7255559) scripts agat_sp_keep_longest_isoform.pl and agat_sp_extract_sequences.pl were used on the fruit fly (*Drosophila melanogaster*) genome assembly (BDGP6.46 (GCA_000001215.4) from Ensembl) and annotation to generate one protein (the longest isoform) per gene. Miniprot [45] was used to align the proteins to the curated assemblies. UniProtKB/Swiss-Prot [46] release 2022_03 in addition to the arthropoda part of OrthoDB v11 [47] were also aligned separately to the assemblies. Red [48] was run via redmask (https://github.com/nextgenusfs/redmask) on the assemblies to mask repetitive areas. In addition, we ran Earl Grey [49] to annotate transposable elements. GALBA [45,50–53] was run with the fruit fly proteins using the miniprot mode on the masked assemblies. The funannotate-runEVM.py script from Funannotate was used to run EvidenceModeler [54] on the alignments of the fruit fly proteins, UniProtKB/Swiss-Prot proteins, arthropoda proteins and the predicted genes from GALBA. The resulting predicted proteins were compared to the protein repeats that Funannotate distributes using DIAMOND blastp and the predicted genes were filtered based on this comparison using AGAT. The filtered proteins were compared to the UniProtKB/Swiss-Prot release 2022_03 using DIAMOND [51] blastp to find gene names and InterProScan was used to discover functional domains. AGAT’s agat_sp_manage_functional_annotation.pl script was used to attach the gene names and functional annotations to the predicted genes. EMBLmyGFF3 [55] was used to combine the fasta files and GFF3 files into a EMBL format for submission to ENA.

### Phylogenetic and comparative genomic analysis

A multi-species comparative analysis was performed using the proteomes from haplotype 1 of the four *Diamesa* species in addition to all the species in Table 1. Peptide sequences and annotations were downloaded from Nell et al. [18] for all species except the *Diamesa* and *Smittia*, while *Smittia* data was taken from Fu et al. [19]. The workflow used was based on the methods from La Torre et al. [56] (https://github.com/RenatoLaTorreRamirez/Algarrobo-Genome). Specifically, for all species, proteins from all chromosomes/scaffolds were used, including only one representative isoform per protein-coding gene in the analyses. Single-copy orthogroups covering all species were inferred using OrthoFinder version 2.5.5 [57], aligned with mafft version 7.515 [58] and trimmed with trimAl version 1.2 [59]. The alignment was used as input for constructing a maximum-likelihood (ML) phylogeny using RAxML version 8.2.12 [60] under the substitution model LG+I+G4+F as determined using ModelTest-NG version 0.1.7 [61]. Phylogenetic hierarchical orthogroups (HOGs) were identified in a second OrthoFinder run specifying the ML tree and then filtered to discard uninformative and large (>100 genes) gene families. This species tree was used along the filtered gene families as inputs for CAFE version 5 [62] to estimate the patterns of expansion or contraction of gene families. A summary and plots of gene family dynamics were obtained using CafePlotter version 0.2.0 (https://github.com/moshi4/CafePlotter).

A gene ontology (GO) enrichment analysis using topGO version 2.59.0 [63] was performed for each of the *Diamesa* species separately, both for significantly contracting and expanding families. A Fisher exact test with the algorithm *weight01* and a *nodeSize* parameter of 10 was used for calculating significance in the terms biological process (BP), molecular function (MF) and cellular component (CC).

### Data availability

The sequencing data, genomes and annotations are available on Zenodo (doi:10.5281/zenodo.15735891).

## RESULTS AND DISCUSSION

### Genome assembly and genome annotation

Here we present the first assembled genomes in the subfamily Diamesinae. Two haplotype-separated genomes could be assembled for all four *Diamesa* species investigated in this study. The final genome assemblies are between 98.9 Mbp and 115.1 Mbp in size, which is within the range of other Chironomidae genomes (Figure 2, Table 1 & Table 2).

**Figure 2:**
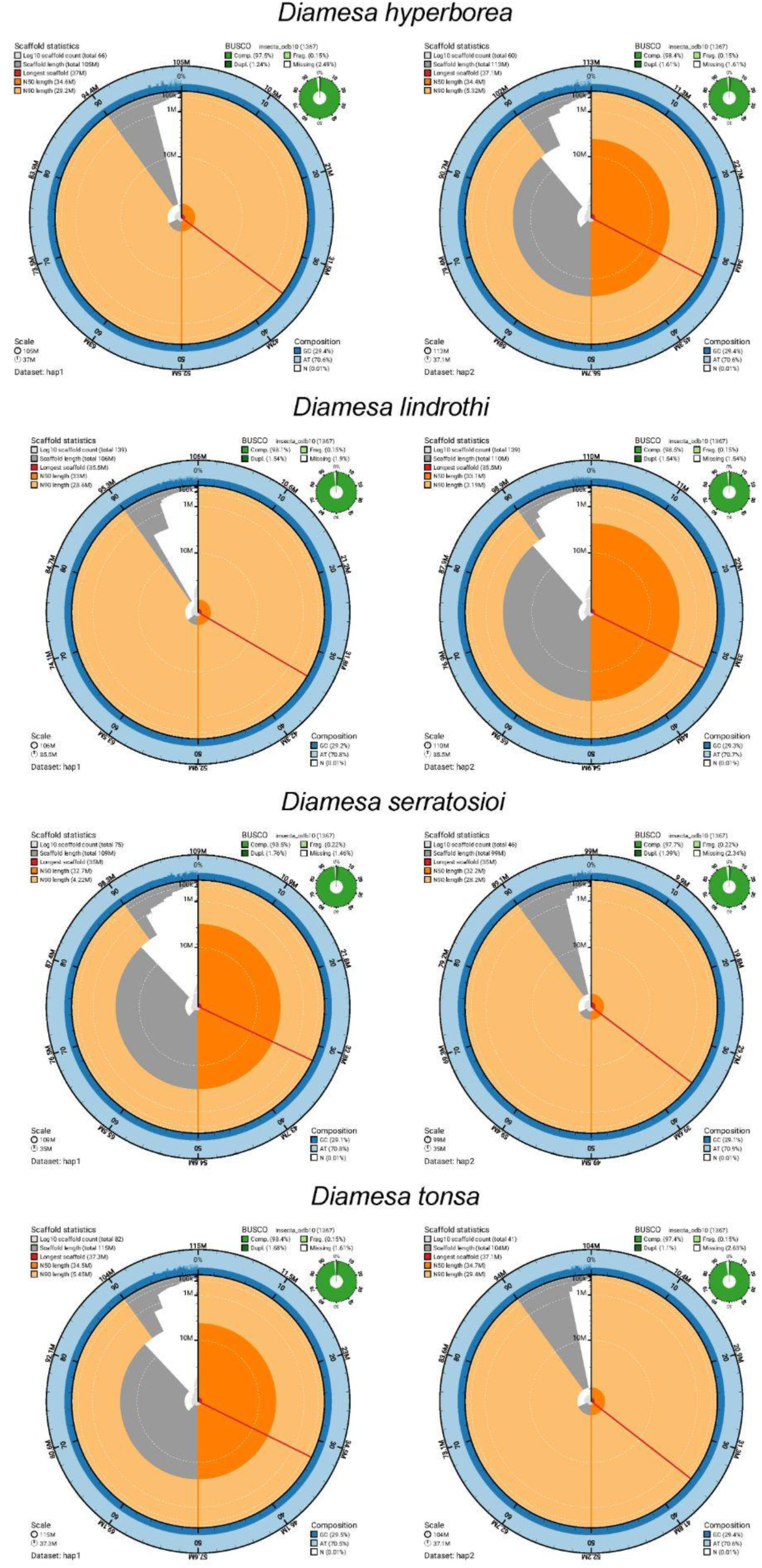
Metrics of the genome assemblies of four *Diamesa* species. The BlobToolKit Snailplots show N50 metrics and BUSCO gene completeness. The two outermost bands of the circle signify GC versus AT composition at 0.1% intervals. Light orange shows the N90 scaffold length, while the deeper orange is N50 scaffold length. The red line shows the size of the largest scaffold. All the scaffolds are arranged in a clockwise manner from the largest to the smallest and are shown in darker gray with white lines at different orders of magnitude. The light gray shows the cumulative scaffold count. The scale inset in the lower left corner shows the total amount of sequence in the whole circle, and the fraction of the circle encompassed in the largest scaffold. For every species, haplotype 1 is depicted on the left, and haplotype 2 is depicted on the right.

**Table 2:** Assembly and annotation statistics for the four *Diamesa* genomes.

| Species | *D. hyperborea* |  | *D. lindrothi* |  | *D. serratosioi* |  | *D. tonsa* |  |
| --- | --- | --- | --- | --- | --- | --- | --- | --- |
| Haplotype | Hap1 | Hap 2 | Hap1 | Hap 2 | Hap1 | Hap 2 | Hap1 | Hap 2 |
| HiFi read coverage | 62x |  | 49x |  | 60x |  | 58x |  |
| Hi-C read coverage | 99x |  | 114x |  | 106x |  | 97x |  |
| Assembly span (Mbp) | 104.9 | 113.4 | 105.8 | 109.9 | 109.2 | 98.9 | 115.1 | 104.5 |
| Number of contigs | 123 | 114 | 189 | 192 | 133 | 98 | 135 | 76 |
| Contig N<sub>50</sub> length (Mbp) | 2.9 | 3.8 | 4.1 | 2.7 | 2.9 | 3.9 | 3.2 | 4.4 |
| Longest contig (Mbp) | 8.8 | 11.3 | 9.1 | 11.3 | 8.9 | 7.8 | 11.1 | 12.5 |
| Number of scaffolds | 66 | 60 | 139 | 139 | 75 | 46 | 82 | 41 |
| Scaffold N50 length (Mbp) | 34.6 | 34.4 | 33.0 | 33.1 | 32.7 | 32.2 | 34.5 | 34.7 |
| Longest scaffold (Mbp) | 36.9 | 37.1 | 35.5 | 35.5 | 35.0 | 34.9 | 37.2 | 37.1 |
| Consensus quality (QV) compared to HiFi (compared to Hi-C) | 64.3<br>(20.9) | 63.5<br>(20.8) | 59.1<br>(30.6) | 59.9<br>(30.9) | 63.1<br>(26.9) | 64.6<br>(26.9) | 63.1<br>(20.7) | 66.7<br>(20.8) |
| Consensus quality (QV) over both haplotypes | 63.9 (20.9) |  | 59.5 (30.8) |  | 63.7 (26.9) |  | 64.5 (20.7) |  |
| K-mer completeness (percentage; compared to HiFi reads) | 73.9 | 77.8 | 91.6 | 92.3 | 89.7 | 86.6 | 74.1 | 70.4 |
| K-mer completeness over both haplotypes | 99.8 |  | 99.8 |  | 99.7 |  | 99.8 |  |
| Genomic BUSCO completeness** | 97.5 | 98.4 | 98.1 | 98.5 | 98.5 | 97.7 | 98.4 | 97.4 |
| Estimated heterozygosity | 1.33% |  | 0.48% |  | 0.64% |  | 1.73% |  |
| **Number of annotated protein-coding genes** | 10798 | 11097 | 10638 | 10831 | 11113 | 10472 | 11375 | 10535 |
| **Number of protein-coding genes with functional domain\*** | 10283 | 10518 | 10181 | 10365 | 10573 | 10037 | 10771 | 10050 |
| **Number of protein-coding genes with gene name** | 6753 | 6842 | 6848 | 6887 | 6944 | 6714 | 6956 | 6680 |
| **Annotation BUSCO completeness\*\*** | 94.9 | 95.1 | 95.7 | 96.1 | 95.9 | 95.1 | 95.5 | 94.8 |

\*Number of genes annotated with a functional domain as found by InterProScan; \*\*Based on the Insecta dataset (v10).

The assembled genomes are slightly larger than their estimated genome size from the *k*-mer spectra (99.8 Mbp, 97.6 Mbp, 92.7 Mbp, and 99.1 Mbp predicted genome size for *D. hyperborea*, *D. lindrothi*, *D. serratosioi*, and *D. tonsa* respectively). Contig and scaffold N_50_ values are generally high, ranging from 2.7-4.4 Mbp for contig N_50_ and 34.9-37.2 Mbp for scaffold N_50_. Three chromosomes were identified in *D. hyperborea*, *D. serratosioi*, and *D. tonsa*, while four chromosomes were identified in *D. lindrothi*. The fourth chromosome in *D. lindrothi* is likely the polytene chromosome typical for chironomid flies. A scaffold of similar size was detected in the three other species, but in each case was not labelled as chromosome due to its presence in only one of the two haplotypes. The presence of four chromosomes is in line with karyotyping in other Diamesinae species [64]. The generated genomes have high BUSCO completeness (>97.4%) (Table 2 and Figure 2), *k*-mer completeness (>99.7% over both haplotypes in all species) (Figure 3), and consensus quality value (QV; >59.1, where a QV of 50 corresponds to one error every 100,000 bp, or 99.999% accuracy). The *k*-mer spectra show a nice separation between *k*-mers derived from sequencing errors (low multiplicity, only found in reads), haplotype-specific *k*-mers (only found in one haplotype, multiplicity around half of the general coverage), and *k*-mers shared between both haplotypes (with a multiplicity similar to the coverage) (Figure 3). Genome annotation of the assemblies identified between 10,472 and 11,375 protein-coding genes (Table 2). In the majority of the proteins (around 95%) at least one functional domain could be detected, and for a large part of the proteins (60-65%), a gene name could be attached. Plots for coverage vs GC (Blobplots) can be found in Supplementary Figure 1, and Hi-C contact maps for the assemblies of the four *Diamesa* species can be found in Supplementary Figure 2.

**Figure 3:**
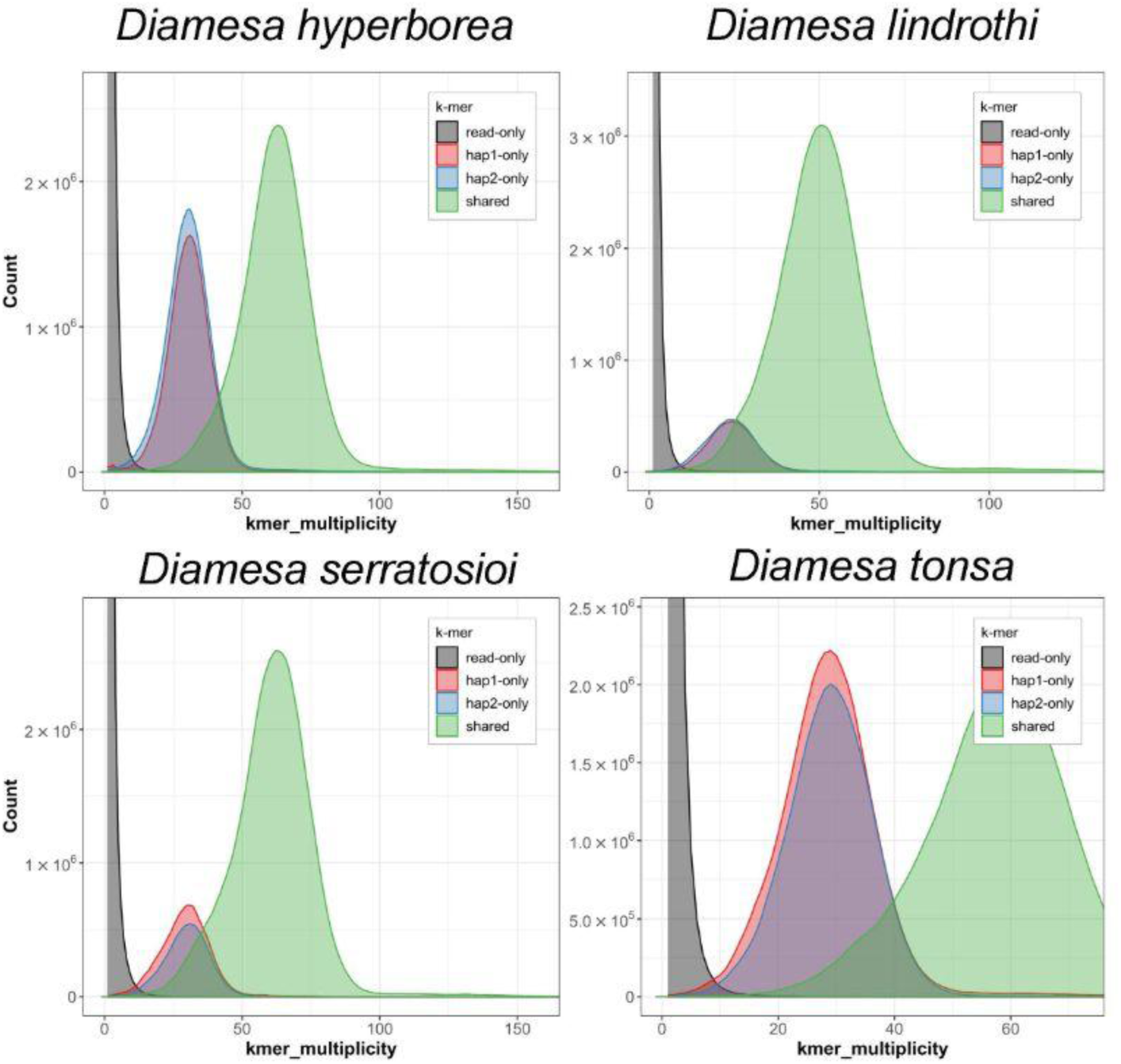
***K*-mer spectra of HiFi reads from the genomes of four *Diamesa* species.** Distributions of *k*-mers found only in the reads (black), only in haplotype 1 (red), only in haplotype 2 (blue), and in both haplotypes (green). The x-axis depicts the number of unique *k*-mers, while the y-axis depicts the *k*-mer multiplicity (how often the *k*-mer is found in the set of HiFi reads).

### Heterozygosity

There was a notable difference in the estimated heterozygosity among the four species sequenced with *Diamesa hyperborea* and *D. tonsa* (Figure 1) showing higher levels than *D. lindrothi* and *D. serratosioi* (Table 2, Figure 3). While more in-depth analyses, preferably with a broader population sampling, would be required to investigate the mechanisms behind this pattern, it aligns with our hypothesis that there is previously occurring or even ongoing introgression between *D. hyperborea* and *D. tonsa*. These two species fly in overlapping time periods at the sample locality in the Rondane Mountains ([65], own observation) and share identical DNA barcodes (dx.doi.org/10.5883/DS-DIANOR1) while harboring considerable differences in their nuclear genomes (see below). A history of interspecific hybridization can explain this pattern.

### Phylogeny

The ML phylogeny based on single-copy orthogroups supports a monophyletic genus *Diamesa* placed as a sister group to the remaining Chironomidae, excepting *Parochlus* (Figure 4). This position, as well as the relationship between other genera and subfamilies in our tree, agree with Cranston et al.’s four-gene phylogeny that had a considerably broader taxon sampling [66]. It is also consistent with the phylogenomic tree presented by Nell et al. [18].

**Figure 4.**
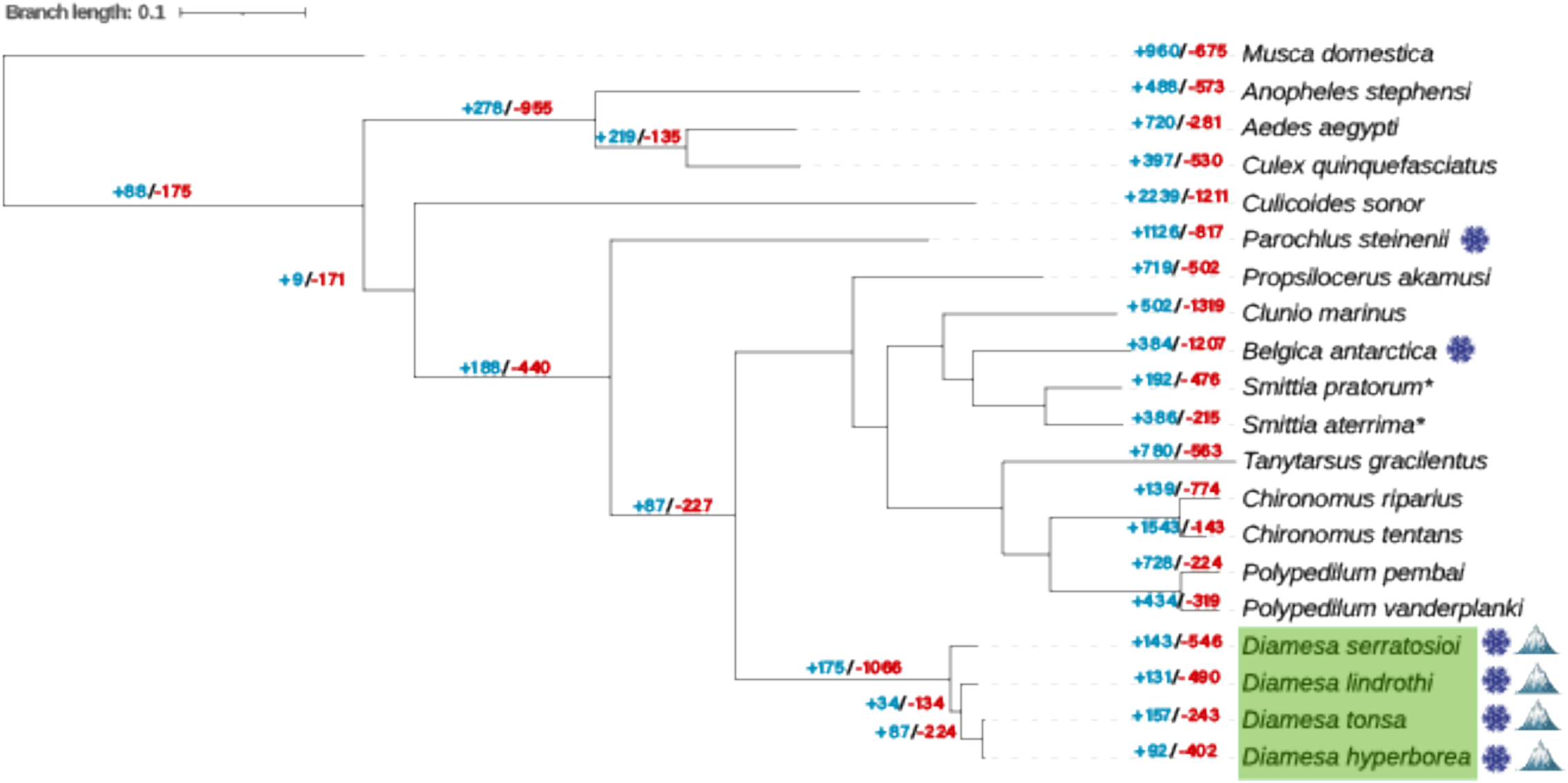
Phylogenomic tree from RAxML and OrthoFinder using single-copy orthogroups rooted with *Musca domestica*. The scale bar represents branch lengths. *Diamesa* are labeled in the green box. A snowflake symbol appears next to species adapted to the cold, while the mountain symbol represents high-altitude adapted species. CAFE analysis was used to determine the number of gene family expansions (in blue) and contractions (in red) for each species and node. *Comparison of COI barcodes indicate that these genomes belong to other species in the genus *Smittia*.

### Comparative genomics

Comparative genomic analysis was used to reveal gene family expansions and contractions within twenty Diptera species’ genomes with the aim of revealing adaptive divergence of the four *Diamesa* species in response to Arctic conditions, namely cold temperatures and high altitudes. CAFE [62] analysis resulted in information about the gene expansions and contractions found for each species and each node (Figure 4). Specifically, we compared primarily the node where our *Diamesa* species diverged from other Diptera (Figure 4). Expanded and contracted gene families significant at the node for *Diamesa* (Figure 4), divided into tables for each of the four *Diamesa* species, can be found in Supplementary Tables 3-10. Results from the GO analysis for the contracted and expanded gene family genes for each of the *Diamesa* species can be found in Supplementary Tables 11 and 12, respectively, and are visualized in Figure 5.

**Figure 5.**
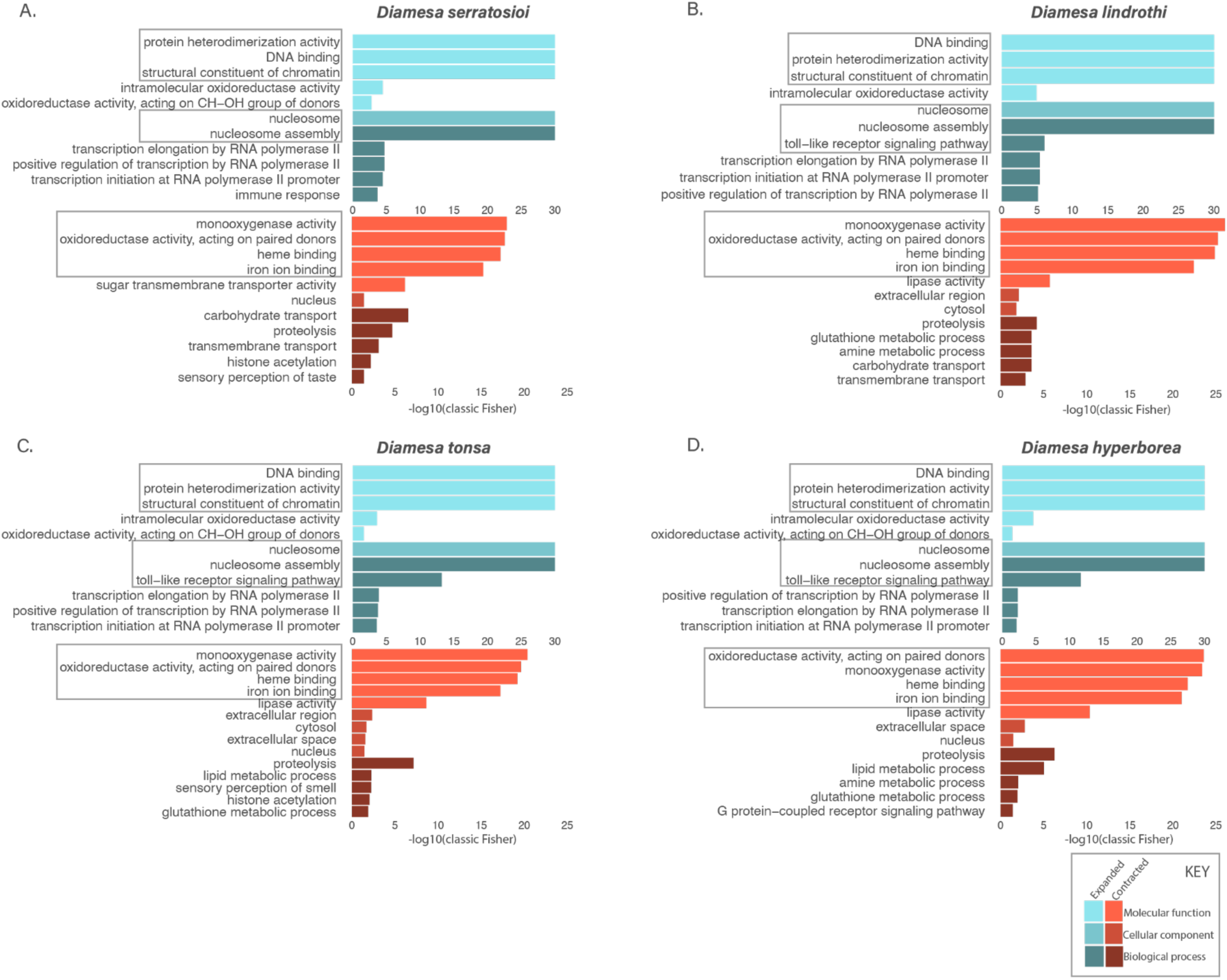
GO analysis of expanded (blue) and contracted (red) gene families. A) *D. serratosioi, B*) *D. lindrothi*, C) *D. tonsa*, and D) *D. hyperborea.* A maximum of the top 5 GO terms per class (biological process, cellular component and molecular function) are presented. The x-axis is plotted as -log10(classic Fisher *p*-value).

### Diamesa tonsa and D. hyperborea are unique species

The large number of differentially expanded and contracted gene families between *D. tonsa* and *D. hyperborea* are strong evidence that these species have undergone divergent evolution and can likely be considered unique species, despite having identical DNA barcodes (dx.doi.org/10.5883/DS-DIANOR1). While not within the scope of this study, with the published genomes of these species, it is now possible to identify regions of divergence between *D. tonsa* and *D. hyperborea* to facilitate more accurate separation of them using molecular markers from the nuclear genome. Such a study should include additional species in the *cinerella*-group such as *D. cinerella* and *D. hamaticornis*, as these have identical or very similar COI DNA barcodes[13].

### Glucose dehydrogenase plays a crucial role in cold-tolerance adaptation

To explore the similarities between the cold-adapted species *B. antarctica, P. steinenii, D. lindrothi, D. serratosio, D. hyperborea* and *D. tonsa*, we searched for gene families with shared contractions of expansions within the genomes of these species. Only one gene family (N0.HOG0000187) was found to be significantly expanded or contracted in the common ancestor of our four *Diamesa* species, as well as *B. antarctica* and *P. steinenii*, while being non-significant in all other species (Supplementary Figure 3). Notably, this gene family significantly expanded in the cold-adapted species was completely absent from *P. vanderplanki*, a desert-dwelling, desiccation-tolerant species, potentially further implicating its importance in cold-adapted species. This gene family includes genes associated with glucose dehydrogenase (GDH) (Uniprot P18173) and ecdysone oxidase (EO) (Uniprot Q9VY01). GDH plays a crucial role in cold adaptation by producing nicotinamide adenine dinucleotide phosphate H (NADPH), which is essential for synthesizing cryoprotectants like sorbitol and glycerol and managing oxidative stress. GDH is involved in the pentose phosphate pathway (PPP), providing NADPH for these processes. Increased GDH activity, and thus NADPH production, in cold-hardy insects may help them survive freezing temperatures by stabilizing cellular structures and reducing damage from reactive oxygen species, in-line with previous reports in the goldenrod gall fly *Eurosta solidaginis* [67,68] and the rice leafroller *Cnaphalocrocis medinalis* [69].

### Genome size as an adaptation to cold and high-altitude

Given that there was only one gene family with an expansion or contraction shared between the cold-adapted species, it is likely that the *Diamesa* have further evolved their own unique genomic mechanisms for cold adaptation. There were 1066 gene family contractions compared to only 175 gene family expansions (Figure 4), representing the largest gene family contraction of any node. Interestingly, the contraction/expansion ratio is similar to the *B. antarctica* branch while showing the opposite trend in *P. steinenii.* Furthermore, given the similarities in genome size for *B. antarctica* (99 Mbp [17]) and the four *Diamesa* species (98-113 Mbp, present study) compared to *P. steinenii* (143 Mbp [23]), could support the hypothesis raised in Kelley et al. [17] suggesting a condensed genome was an evolutionary mechanism for cold-tolerance. Although clearly this indicates it is not the primary evolutionary strategy for overcoming freezing temperatures, as is the case with *P. steinenii* having a substantially larger genome. To investigate similarities between these small genome cold-adapted Diptera species, we investigated shared gene family contractions/expansions between only *B. antarctica* and the four *Diamesa*. Surprisingly, this resulted in no significantly contracted/expanded gene families in common that were unique to these species, indicating that small genome size could be related to a lack of repeats, TEs and introns rather than to the dynamism of particular gene families. In fact, the correlation between genome size drivers in Chironomidae was further explored in Nell et al. [18], where the authors determined the small genome sizes in this family was as a result of loss of noncoding regions and repeat elements, although *Diamesa* was not included in their analysis. To better understand the evolution of the small genome size of *Diamesa* would require a detailed comparative analysis of all available chironomid genomes and additional Diamesinae species are required to determine genomic characteristics, such as the number of TEs and repeats, as well as intron lengths, intergenic site lengths and recombination rate across genomes [5,70,71].

### Histone and toll protein gene family expansions in Diamesa

Given the lack of similarity in shared gene family dynamics between our *Diamesa* species and other cold-tolerant species in this study, we next focused on the gene contractions and expansions that were unique to the *Diamesa* node. Of the gene families reported in the *Diamesa* node, 88 and 13 were significantly contracted and expanded, respectively. These gene families were further investigated individually, and used to determine enriched GO terms from each list (Supplementary Tables 10 and 11).

The expanded gene families were predominantly related to histones and toll proteins, and to a lesser extent with GDH (discussed above) (Supplementary Tables 3-6). This was further reflected in the GO analysis of the expanded gene families, which was highly enriched for terms involving chromatin/histone/nucleosome (i.e. GO:0030527/GO:0000786/GO:0006334) as well as Toll-like receptor (TLR) proteins (GO:0002224) (Figure 5, Supplementary Table 12). The interplay between the highly enriched nucleosome term with the expanded gene families associated with histones can be explained by their roles in epigenetic gene regulation. In insects, post-translational histone modifications have been linked to the lifespan and energy utilization of numerous insects, including in the *Eurosta solidaginis* and the goldenrod gall moth *Epiblema scudderiana* under zero-temperature conditions [72], and even in diapausing mosquitoes [73].

The TLR pathway, an ancient regulatory cascade involved in host defense [74], is essential to the innate immune system in insects, combating fungal and bacterial pathogens. Recent research indicates that environmental conditions, such as cold stress, can influence immune function. In *Drosophila*, cold-induced immune activation is hypothesized to compensate for reduced immune efficiency at low temperatures. Furthermore, environmental temperature has been demonstrated to significantly alter immune responses and the energetic costs of immunity in larvae of the yellow mealworm beetle *Tenebrio molitor*. Sinclair et al. [75] reviewed tolerance mechanisms in insects to cold, and argued that there is a relationship between low temperatures and the immune response in insects. Thus the expansion of this gene family in our cold adapted species further supports this hypothesis.

A study by Kim et al. [76] similarly noted expansions in histone-related gene families as well as GO term enrichment for regulation of the TLR signaling pathway in *B. antarctica* and *P. steinenii*, further highlighting the necessity of these mechanisms for cold adaptation. However, in a more recent comparison of chironomid midges by Nell et al. [18], their analysis did not indicate an expansion in gene families associated with histones, chromatin, nor TLR signaling pathways, indicating that these expansions occurring in *Diamesa* could have evolved concertedly in cold-tolerant Chironomidae lineages. Further investigations into the role of the expanded histone-related gene families could involve gene expression analyses in *Diamesa,* as well as other cold-tolerant Diptera, at varying temperatures to determine if and which genes are having altered gene expression due to these histone modifications.

### Contractions in gene families associated with oxygen transport and metabolism in Diamesa

As mentioned above, there were substantially more gene families undergoing contractions than expansions. This preference for *Diamesa* gene families to contract is likely a result of selective pressures due to rapid changes in their environmental conditions. This is in line with previous findings of gene loss, rather than gene function, contributing to the adaptive evolution of a variety of organisms, from yeast [77] and bacteria [78], to mammals [79] and insects [80]. In a recent study of convergent evolution of high-altitude adapted mammals [79], the authors found that the convergence of the gene family contractions in high-altitude species is much greater than that of expansion, with many of these gene families related to hypoxia response. This study of mammals highlights, in conjunction with our findings in insects, a potential cross-kingdom mechanism of high-altitude animal adaptation that warrants further study. To fully explore this hypothesis, comparative genomic studies are needed in a diversity of high-altitude adapted Diptera species.

More specifically, of the gene annotations within the contracted gene families (Supplementary Tables 7-10), the terms appeared diverse, as opposed to the clear histone signal in the expanded genes (Supplementary Table 3-6). However, after performing GO term enrichment analysis (Figure 5, Supplementary Table 11), we were able to determine that most significantly enriched GO terms, incorporating oxidoreductase activity, monooxygenase activity, heme binding and iron ion binding, are related to the function of heme-containing proteins, particularly those involved in oxygen transport and metabolism.

High-altitude animals commonly exhibit positive selection and rapid evolution of genes involved in hypoxia response, suggesting that general genetic mechanisms might be utilized to adapt to high-altitude extremes [81,82]. Furthermore, changes in oxygen levels in an organism’s environment can drive natural selection for or against genes involved in oxygen metabolism. Previous studies on the convergent evolution of different species, including mammals, fungal pathogens and insects, also indicated that when species face selection pressure such as environment or eating habits, the expansion and contraction of gene families are mainly the result of direction changes [80,83]. As such, the contraction of gene families associated with oxygen transport and metabolism could be interpreted as these processes being either specialized within these northern *Diamesa* allowing for a reduction of a larger array of genes controlling these functions, or could indicate less demand of oxygen metabolism in response to their environmental needs, i.e. high altitude and cold temperatures.

While there are no studies reporting contractions of gene families in oxygen transport/metabolism in high-altitude Diptera, a striking overlap between GO terms downregulated in cold-acclimated *D. melanogaster* [84] and *Diamesa* contracted gene families was noted (Figure 5). This included GO terms associated with oxidoreductase activity (GO:0016705), heme binding (GO:0020037), iron ion binding (GO:0005506), lipase activity (GO:0016298), extracellular space (GO:0005615) and extracellular region (GO:0005576), proteolysis (GO:0006508), and lipid metabolic processes (GO:0006629). While *Drosophila* and *Diamesa* are not closely related, they both belong to the order Diptera, indicating that these oxygen metabolism processes were already under selection in the *Drosophila* lineage’s response to cold and further evolved as gene redundancies in *Diamesa*.

## CONCLUSIONS

This comparative genomic study reveals significant insights into the adaptive mechanisms of Arctic *Diamesa* species. The marked contraction of gene families involved in oxygen transport and metabolism suggests a potential reduction in reliance on certain hypoxia-related processes, possibly reflecting adaptations to high-altitude and cold environments where oxygen demand may be decreased or where oxygen metabolism processes are highly specialized. Conversely, the expansion of gene families related to histones and TLR signaling pathways indicates a strategic enhancement of gene regulation and immune responses, likely crucial for survival under extreme cold stress. The notable expansion of glucose dehydrogenase underscores its role in cold tolerance through cryoprotectant synthesis and oxidative stress management. Collectively, these genomic features highlight the complex interplay of gene family contraction and expansion that underpin the unique cold adaptation strategies of *Diamesa* species. Further functional and gene expression studies could elucidate these mechanisms and confirm their roles in ecological resilience to the harsh high-latitude environments.

Furthermore, the generation of four haplotype-resolved reference genomes for these non-model *Diamesa* species represents a valuable resource for future research. Such high-quality, phased genomes provide a more accurate representation of genetic diversity and facilitate detailed analyses of adaptive variation, gene flow, and evolutionary history. They are essential for identifying structural variants, understanding heterozygosity, and elucidating the genetic basis of traits associated with extreme environments. This genomic resource thus lays a foundation for ongoing studies in ecological genomics, conservation, and evolutionary biology of Arctic and high-altitude insects.

## ACKNOWLEDGMENTS

Thanks to the County Governour of Innlandet and Rondane-Dovre Nasjonalparkstyre for permission (022/5001-4 432.3) to collect chironomids within the borders of Rondane National Park. Generation of the genomes in this study were performed as part of Research Council of Norway project 326819 (The Earth Biogenome Project Norway; EBP-Nor). This project received data management and infrastructure support from ELIXIR Norway, supported by the Research Council of Norway’s grant 270068, the University of Bergen, the University of Oslo, the Arctic University of Norway in Tromsø, the Norwegian University of Science and Technology and the Norwegian University of Life Sciences. This study was also supported in part by the Norwegian Directorate for Higher Education and Skills, through the Norwegian Partnership Programme for Global Academic Cooperation (NORPART), project NORPART2021/10475 ‘BiGTREE’. The authors acknowledge support from the National Infrastructure for High Performance Computing and resources provided by Sigma2 as well as Data Storage in Norway (project NN8013K) for computational work. The Norwegian Sequencing Centre generated the sequencing data used in this project (http://sequencing.uio.no).

