## Supplementary Figure for "Haplotype-resolved chromosome-level genome assemblies of four *Diamesa* species reveal the genetic basis of cold tolerance and high-altitude adaptations in arctic chironomids"

**TITLE**


**AUTHORS & AFFILIATIONS**

Sarah L.F. Martin[1*], Renato La Torre[1], Bram Danneels[2], Ave Tooming-Klunderud[3], Morten Skage[3], Spyridon Kollias[3], Ole Kristian Tørresen[3], Mohsen Falahati Anbaran[1], Elisabeth Stur[1], Kjetill S. Jakobsen[3,] Michael D. Martin[1#], Torbjørn Ekrem[1#*]

[1]Department of Natural History, NTNU University Museum, Norwegian University for Science and Technology, NO-7491 Trondheim, Norway
[2]Computational Biology Unit, Department of Informatics, University of Bergen, Norway
[3]Centre for Ecological and Evolutionary Synthesis, Department of Biosciences, University of Oslo, Norway

[#]Indicates shared senior authorship

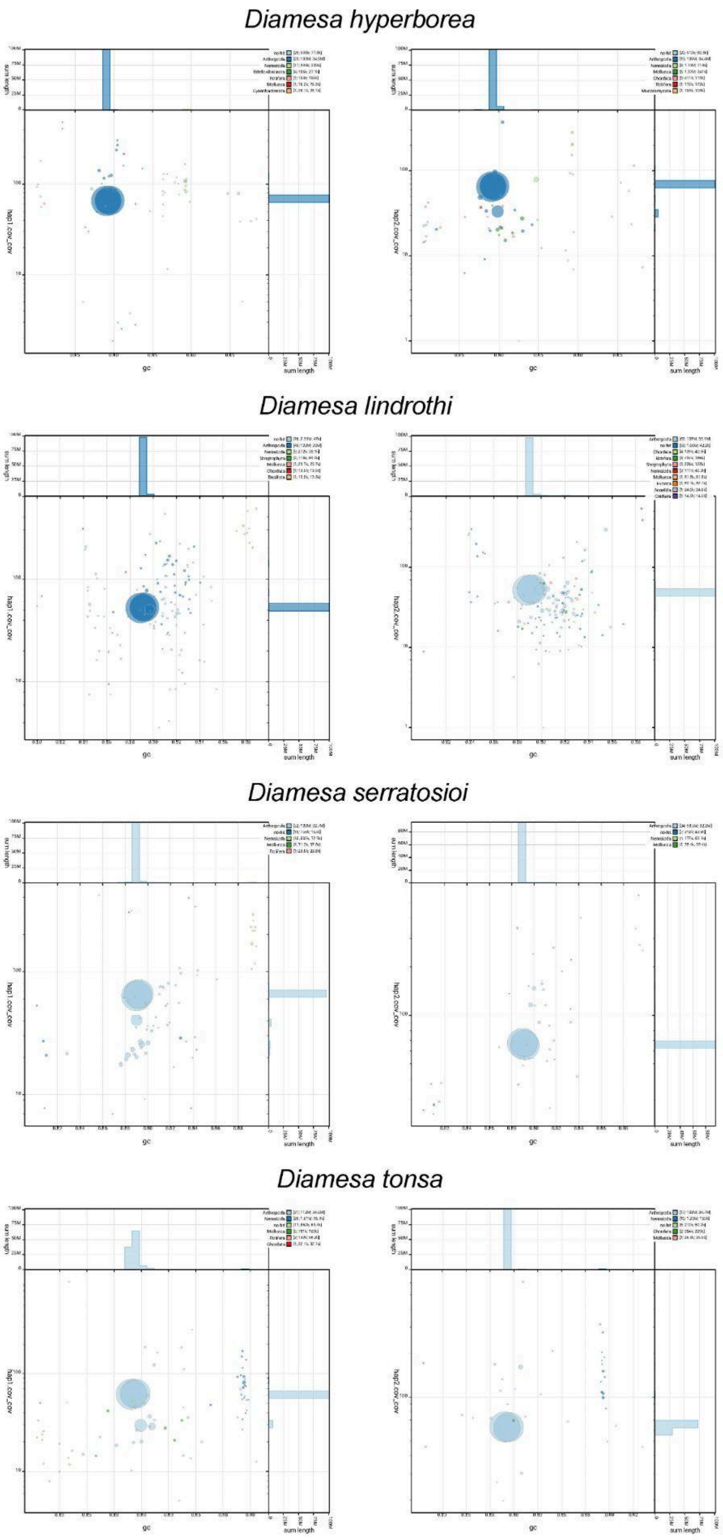


**Supplementary Figure 1: Coverage vs GC plots of the four *Diamesa* species.** The BlobToolKit Blobplots depicts each scaffold as a dot based on the GC content (%GC, x-axis) and coverage (Y-axis). Size of the dots correspond to scaffold length. Dots are colored based on assigned taxonomy. Histograms of sequence lengths within a certain %GC range or coverage range are depicted on the top and right respectively.

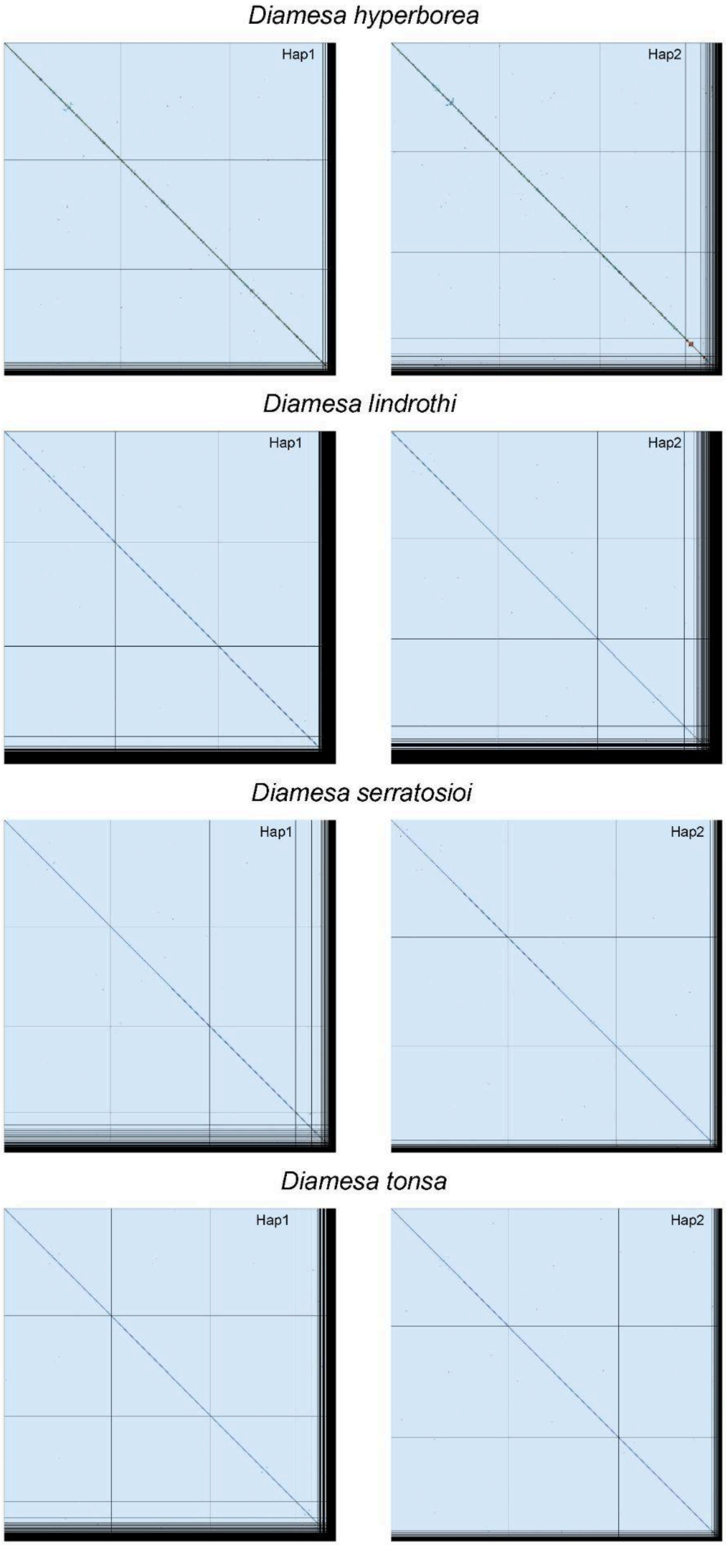


**Supplementary Figure 2**: **Hi-C contact map for the assemblies of the four *Diamesa* species.** The contact map displays interaction frequencies between genomic regions, where darker shades represent a higher number of Hi-C contacts. The axes correspond to the coordinates along each assembly. Hi-C contact maps were generated using PretextMap and visualized using PretextSnapshot.

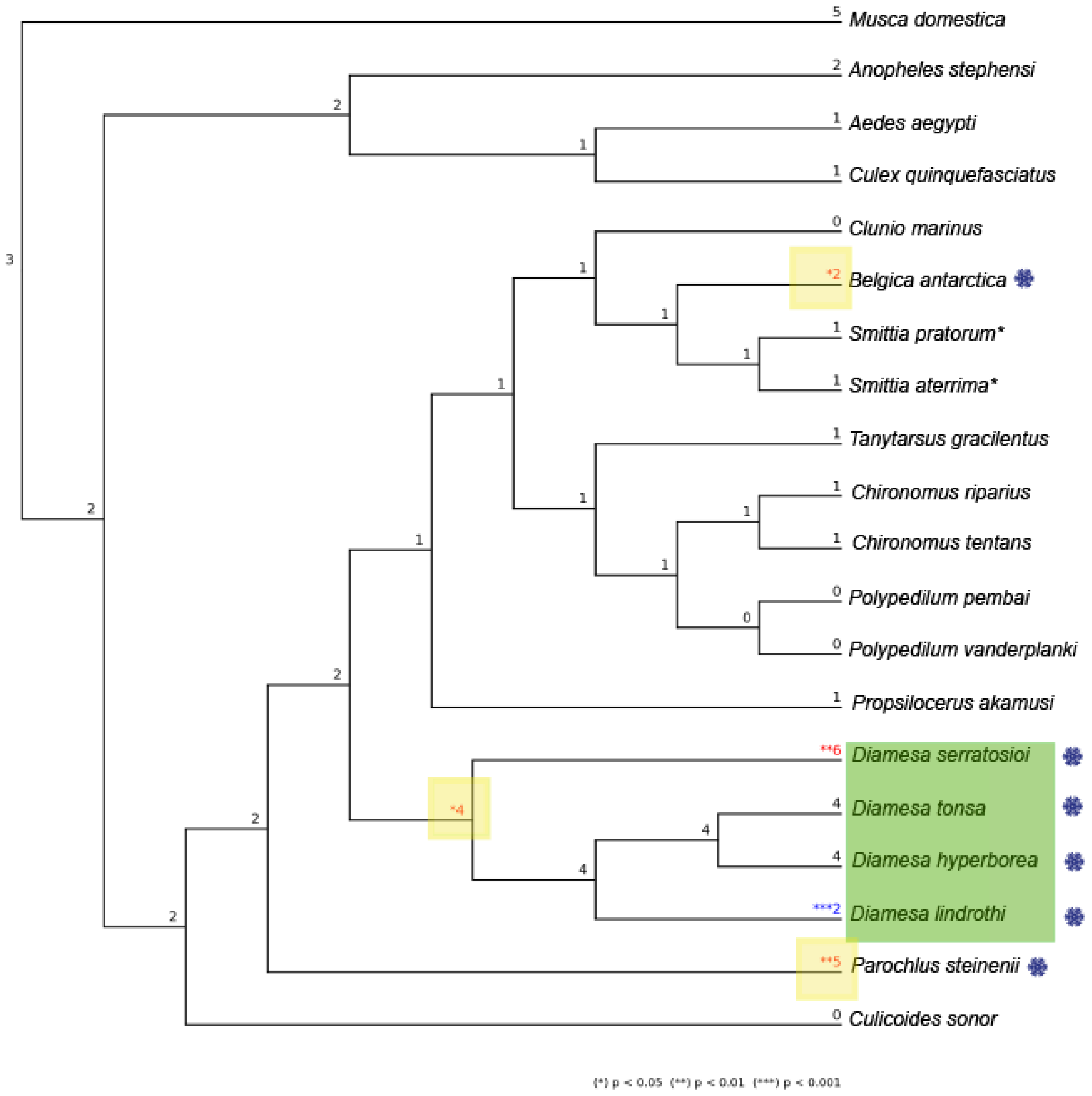


**Supplementary Figure 3**: **Tree output from OrthoFinder showing the number of significant expansions (red) and contractions (blue) for the gene family N0.HOG0000187**. Significance of $p < 0.05$ is noted with *, $p < 0.01$ is noted with **, and $p < 0.001$ is noted with ***. The green box indicates the *Diamesa* species, the yellow boxes indicate the nodes/branches with significant expansions of this gene family. The snowflakes indicate the cold adapted species.
